# Evidence for loss and adaptive reacquisition of alcoholic fermentation in an early-derived fructophilic yeast lineage

**DOI:** 10.1101/213686

**Authors:** Carla Gonçalves, Jennifer H. Wisecaver, Madalena Salema-Oom, Maria José Leandro, Xing-Xing Shen, David Peris, Chris Todd Hittinger, Antonis Rokas, Paula Gonçalves

## Abstract

Fructophily is a rare trait that consists in the preference for fructose over other carbon sources. Here we show that in a yeast lineage (the *Wickerhamiella/Starmerella*, W/S clade) formed by fructophilic species thriving in the floral niche, the acquisition of fructophily is part of a wider process of adaptation of central carbon metabolism to the high sugar environment. Coupling comparative genomics with biochemical and genetic approaches, we show that the alcoholic fermentation pathway was profoundly remodeled in the W/S clade, as genes required for alcoholic fermentation were lost and subsequently re-acquired from bacteria through horizontal gene transfer. We further show that the reinstated fermentative pathway is functional and that an enzyme required for sucrose assimilation is also of bacterial origin, reinforcing the adaptive nature of the genetic novelties identified in the W/S clade. This work shows how even central carbon metabolism can be remodeled by a surge of HGT events.

## Introduction

Comparative genomics is a powerful tool for discovering links between phenotypes and genotypes within an evolutionary framework. While extraordinary progress in this respect has been observed in all domains of life, analyses of the rapidly increasing number of fungal genomes available has been particularly useful to highlight important aspects of eukaryotic genomes, including a broader scope of evolutionary mechanisms than was thus far deemed likely. For example, horizontal gene transfers (HGT), are thought to have played a very important role in domestication (Gibbons et al., 2012; Marsit et al., 2015; Ropars et al., 2015) and in the evolution of metabolism in fungi (Alexander, Wisecaver, Rokas, & Hittinger, 2016; Wisecaver & Rokas, 2015). Instances of the latter are best showcased by the high frequency of HGT events involving gene clusters related to fungal secondary metabolism (Campbell, Rokas, & Slot, 2012; Khaldi & Wolfe, 2011; Wisecaver & Rokas, 2015). When considering the horizontal transfer of single genes, those encoding nutrient transporters seem to be among the most frequently transferred (Coelho, Goncalves, Sampaio, & Goncalves, 2013; Goncalves, Coelho, Salema-Oom, & Goncalves, 2016; T. A. Richards, 2011). While HGT events are reasonably easy to detect in a reliable manner given sufficient sampling of the lineages under study, inferences concerning the evolutionary driving forces behind HGT are often difficult and uncertain, because most HGT events identified are ancient. However, available evidence suggests that HGTs are often associated with rapid adaptation to new environments (Cheeseman et al., 2014; Gojkovic et al., 2004; Qiu et al., 2013; Thomas A. Richards et al., 2011; Thomas A. Richards & Talbot, 2013).

In line with these findings, we recently reported on the evolutionary history of a unique, high-capacity, specific fructose transporter, Ffz1, which is intimately associated with fructophilic metabolism in ascomycetous yeasts (Saccharomycotina) (Goncalves et al., 2016). Fructophily is a relatively rare trait that consists in the preference for fructose over other carbon sources, including glucose (Cabral, Prista, Loureiro-Dias, & Leandro, 2015; Goncalves et al., 2016; Sousa-Dias, Gonçalves, Leyva, Peinado, & Loureiro-Dias, 1996). The evolution of *FFZ1* involved the likely horizontal acquisition of the gene from filamentous fungi (Pezizomycotina) by the most recent common ancestor (MRCA) of an early-derived lineage in the Saccharomycotina, composed so far entirely of fructophilic yeasts (Goncalves et al., 2016). Most of the approximately one hundred species forming this clade (*Wickerhamiella* and *Starmerella* genera, as well as closely related *Candida* species), are associated with the floral niche and are often isolated from fructose-rich nectar (Canto, Herrera, & Rodriguez, 2017; de Vega et al., 2017; Lachance et al., 2001). Interestingly, fructophilic lactic acid bacteria, whose metabolism has been dissected in detail, also populate the floral niche (Endo, Futagawa-Endo, & Dicks, 2009; Endo & Salminen, 2013). These bacteria have been shown to grow poorly on glucose, which can be at least partly explained by their lack of respiratory chain enzymes and alcohol dehydrogenase activity, deficiencies that hinder NAD^+^ regeneration during growth on this sugar, as shown for *Lactobacillus kunkei* (Maeno et al., 2016). In contrast to glucose, fructose can be used both as a carbon source and as an electron acceptor for the re-oxidation of NAD(P)H (Zaunmuller, Eichert, Richter, & Unden, 2006), providing an explanation on why it is favored over glucose. Hence, fructophily in lactic acid bacteria seems to be linked to redox homeostasis (Endo, Tanaka, Oikawa, Okada, & Dicks, 2014). In yeasts, it is still unclear how preferential consumption of fructose may be beneficial, partly because unlike fructophilic bacteria, fructophilic yeasts grow vigorously on glucose when it is the only carbon and energy source available (Sousa-Dias et al., 1996; Tofalo et al., 2012). Our previous work suggested that, although a strict correlation was found so far between the presence of Ffz1 and fructophily in all species investigated (Cabral et al., 2015; Goncalves et al., 2016; Leandro, Cabral, Prista, Loureiro-Dias, & Sychrova, 2014) and the requirement for *FFZ1* was genetically confirmed in the fructophilic species *Zygosaccharomyces rouxii* (Leandro et al., 2014), it is very likely that there are additional requirements for fructophily. Thus, the *FFZ1* gene does not seem to be sufficient to impart a fructophilic character to a previously glucophilic species.

To gain insight into the genetic underpinnings of fructophily in yeasts and how it may have become evolutionarily advantageous, here we used comparative genomics to identify traits, other than the presence of the *FFZ1* gene that might differentiate yeasts in the fructophilic *Wickerhamiella/Starmerella* (W/S) clade, focusing on central carbon metabolism. Our results suggest that the evolution of fructophily may have been part of a process of adaptation to sugar-rich environments, which included a profound remodeling of alcoholic fermentation involving the acquisition of bacterial alcohol dehydrogenases and an invertase, the latter being essential for sucrose assimilation. In general, we found a higher than expected number of HGT events in the W/S clade when compared with other lineages in the Saccharomycotina (Marcet-Houben & Gabaldon, 2010) and that a significant number of genes horizontally acquired from bacteria seem to impact redox homeostasis.

## Results

### The horizontally transferred Ffz1 transporter is essential for fructophily in *St. bombicola*

We previously reported the acquisition of a high-capacity fructose transporter (Ffz1) through HGT by the MRCA of W/S-clade species. This transporter was lost in the MRCA of the Saccharomycotina and was later transferred from a Pezizomycotina-related species to the MRCA of the W/S clade, and then from the W/S clade to the MRCA of the *Zygosaccharomyces* genus (Goncalves et al., 2016). A putative role for Ffz1 in fructophily in the W/S clade was hypothesized based on its kinetic properties (Pina, Goncalves, Prista, & Loureiro-Dias, 2004) and the evidence that it is indispensable for fructophily in the phylogenetically distant species *Z. rouxii* (Leandro et al., 2014). To test this hypothesis, a *FFZ1* deletion mutant was constructed in the genetically tractable W/S-clade species *St. bombicola*. The sugar-consumption profile in YP medium supplemented with 10% (w/v) fructose and 10% (w/v) glucose (conditions where fructophily is apparent, hereafter referred to as 20FG medium), showed that fructophilic behavior was completely abolished in the *ffz1∆* mutant (Figure 1), similarly to what was found in *Z. rouxii* (Leandro et al., 2014). A slight increase in the glucose consumption rate was also observed for the *ffz1∆* mutant compared to the Wild Type (WT) when cultures were grown in 20FG medium (Figure 1).

**Figure 1.**
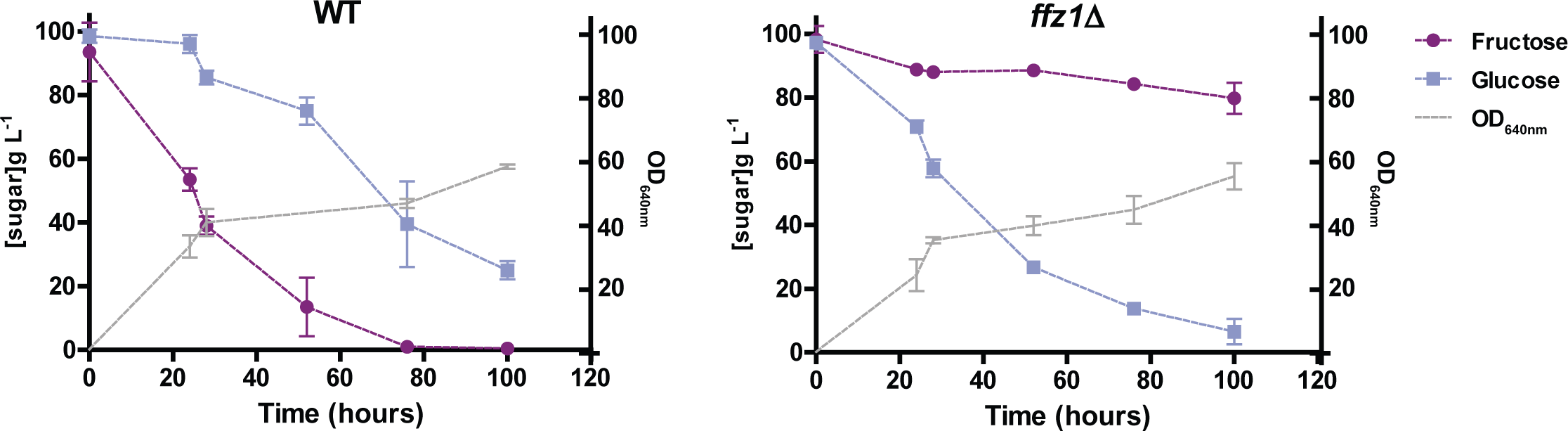
Sugar consumption profiles of *St. bombicola* Wild Type (WT) and *ffz1Δ*. Strains were grown in YP supplemented with 10% (w/v) fructose and 10% (w/v) glucose for 100 hours at 30ºC with aeration. Error bars represent standard deviation of assays performed in duplicate in two biological replicates.

### Comparative genomics of sugar metabolism in W/S clade yeasts

The presence of the *FFZ1* gene was so far the only genetic trait identified to be common to all W/S-clade species investigated that distinguished this clade from its neighboring species in the Saccharomycotina phylogeny (Goncalves et al., 2016). In order to search for other possible signs of the adaptation of sugar metabolism in W/S-clade genomes, a tBLASTx search using sugar metabolism-related genes as queries (Figure 2-source data 1) was performed on a custom database containing the draft genomes of W/S-clade species. The draft genome of *Z. kombuchaensis*, which belongs to the only other Saccharomycotina clade comprising fructophilic yeasts, and publicly available genomes of the closest relatives of W/S-clade species (*Sugiyamaella lignohabitans* and *Blastobotrys adeninivorans*) were also included in the database. For all species, except those comprising the W/S clade, all genes identified in this survey were found to encode proteins exhibiting the expected degree of similarity to other yeast homologs (Figure 2-source data 1). In contrast, in W/S-clade genomes, while all the essential enzymes forming the pentose phosphate pathway and glycolysis presented the expected level of similarity to other yeast orthologs, tBLASTx results for the two genes required for the conversion of pyruvate into ethanol (de Smidt, du Preez, & Albertyn, 2008; Flikweert, van Dijken, & Pronk, 1997) yielded a markedly different result (Figure 2-source data 1). In these species, *PDC1* (encoding pyruvate decarboxylase) and *ADH1* (encoding the main alcohol dehydrogenase), had a lower sequence identity to *S. cerevisiae* homologs than anticipated (~40% in both cases, E-value >e^−80^). This is particularly evident in the comparison with the tBLASTx results obtained for the species most closely related to the W/S clade, *Su. lignohabitans* and *B. adeninivorans*, for which identity of Adh1 and Pdc1 to *S. cerevisiae* homologs is around 60–70% (E-value < e^−130^). A similar result (i.e. at odds with the expected level of similarity) was obtained for the gene encoding invertase (*SUC2*), which is essential for sucrose assimilation in *S. cerevisiae* (Carlson, Osmond, & Botstein, 1981; Gascon, Neumann, & Lampen, 1968).

**Figure 2.**
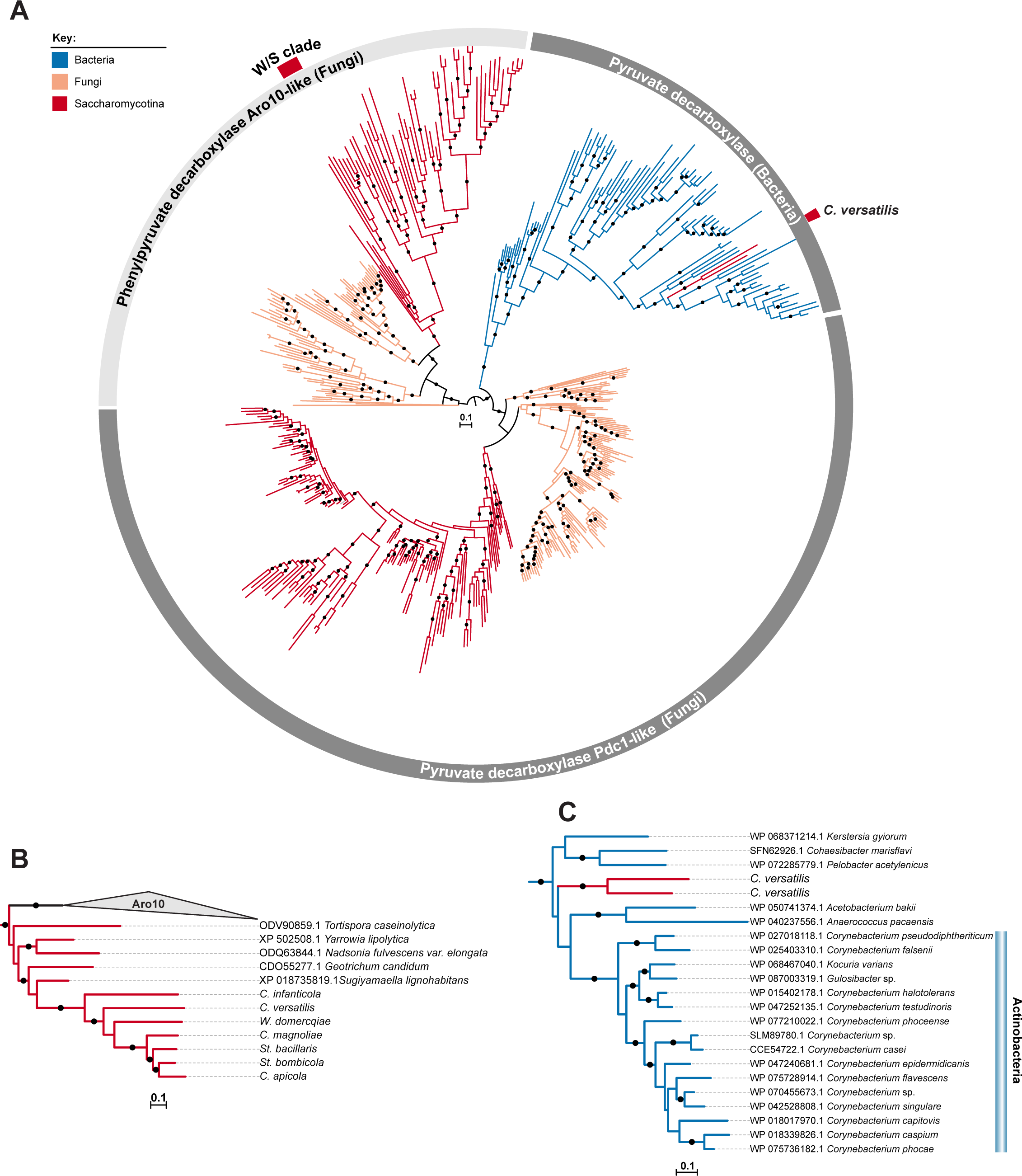
ML phylogeny of Pdc1 and Aro10. A) ML phylogeny of Pdc1-like proteins. W/S-clade and *C. versatilis* sequences are indicated by red blocks. Only branches with bootstrap support higher than 90% are indicated (black dots). Clades highlighted in grey (Aro10- and Pdc1-like) were assigned according to the phylogenetic position of functionally characterized *S. cerevisiae* proteins. B, C) Pruned ML phylogenies depicting the phylogenetic relationship between W/S-clade Aro10 proteins and their closest relatives in the Saccharomycotina (B) and between *C. versatilis* Pdc1 xenologs and the closest related bacterial pyruvate decarboxylases (C).

To better understand the results observed for the *ADH1*, *PDC1*, and *SUC2* genes present in W/S clade genomes, a subsequent BLASTp search against the National Center for Biotechnology Information (NCBI) non-redundant database was performed using the respective W/S-clade best hits for each of the three genes. While W/S clade Adh1 and Suc2 proteins showed higher identity to bacterial proteins (~60%) than to yeast homologs (~45%), the Pdc1-like sequence showed high sequence identity to those of closely related decarboxylases from the Aro10 family (best NCBI hits were Aro10-like proteins from *Su. lignohabitans* and *Geotrichum candidum*). In *S. cerevisiae*, *ARO10* encodes a decarboxylase that is not involved in alcoholic fermentation. Surprisingly, two additional Pdc1 hits were obtained for the *C. versatilis* proteome (Figure 2- source data 1). A subsequent BLASTp search against the NCBI database, confirmed that these two sequences have higher identity to bacterial pyruvate decarboxylases (~44%) than to Saccharomycotina homologs (~35%). To clarify the origin of the Pdc1-like proteins found in W/S genomes, a Maximum Likelihood (ML) phylogeny was constructed using the top 500 NCBI BLASTp hits using *S. cerevisiae* Pdc1 (CAA97573.1), *St. bombicola* Pdc1/Aro10-like and *C. versatilis* Pdc1-like sequences from apparent bacterial origin as queries. Pdc1/Aro10-like sequences from the other W/S-clade species not available at the NCBI database were also included. This phylogeny (Figure 2A) confirmed clustering of the W/S-clade Aro10-like sequences with other Aro10 proteins from fungi, which implies that *PDC1* was lost in the W/S clade. Additionally, as suggested by the BLASTp results, the two Pdc1-like proteins from *C. versatilis* were clustered with bacterial pyruvate decarboxylases (Figure 2C).

### Identification of genes of bacterial origin in the W/S clade

The tBLASTx analyses uncovered genes relevant for sugar metabolism in the genomes of W/S clade yeasts that were possibly horizontally transferred from bacteria (*ADH1*, *PDC1*, and *SUC2*). To identify other genes that fell outside this narrow scope, a systematic high-throughput analytical pipeline based on the Alien Index (AI) score (Alexander et al., 2016; Gladyshev, Meselson, & Arkhipova, 2008) was employed for HGT-detection. In this analysis, we included six W/S-clade species and one additional species, *Candida infanticola* (Kurtzman, 2007), that is phylogenetically positioned as an outgroup of the W/S clade (Figure 2-figure supplement 1), lacks the Ffz1 transporter, and does not inhabit sugar-rich habitats. After implementation of both AI (AI > 0.1) and alignment thresholds, 52 ortholog groups were found with representatives in two or more W/S-clade genomes. As expected, *ADH1* and *SUC2* were present in this group (Figure 3-source data 1). *PDC1* was not detected using these thresholds because only ortholog groups of bacterial origin detected in at least two W/S-clade species were selected, and bacterial Pdc1 proteins were only identified in *C. versatilis*.

**Figure 3.**
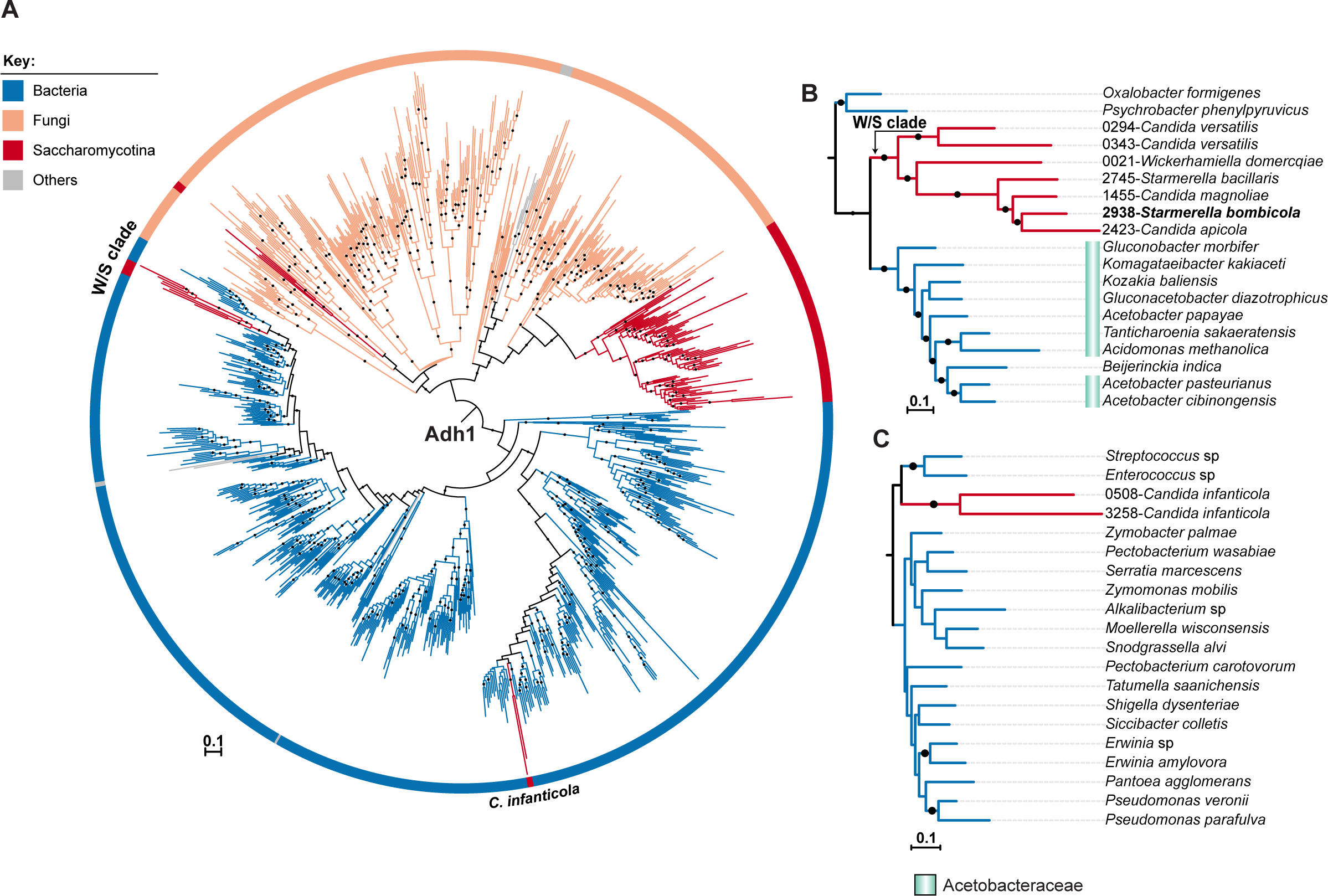
ML phylogeny of fungal and bacterial Adh1 proteins. A) ML phylogeny of Adh1 proteins (top 4.000 phmmer hits). W/S-clade species and *C. infanticola* are highlighted. Only branches with bootstrap support higher than 90% are indicated (black dots). Poorly represented lineages (<10 sequences) are shown in grey. B, C) Pruned ML phylogenies of Adh1 depicting the phylogenetic relationship between the W/S clade and Acetobacteraceae (B) and between *C. infanticola* and other groups of bacteria (C).

To check for putative enrichment in particular protein functions among transferred genes, the 52 candidates were cross-referenced with Kyoto Encyclopedia of Genes and Genomes (KEGG), Gene Ontology (GO), and InterPro annotations. Notably, out of 43 proteins to which a GO molecular function was assigned, 16 impacted redox homeostasis and were associated with oxidoreductase activity (GO:0016491 and GO:0016616, 14 genes) and peroxidase activity (GO:004601, two genes), while a BLAST KOALA annotation (Kanehisa, Sato, & Morishima, 2016) indicated that the biological processes most frequently involved were amino acid metabolism, carbohydrate metabolism, and metabolism of cofactors and vitamins (Figure 3- source data 1). Interestingly, some of these genes appeared to have undergone several intraspecies duplications, in particular those encoding dehydrogenases that participate in various metabolic pathways.

### HGT and gene duplication of alcohol dehydrogenase genes in W/S-clade species

Interestingly, in addition to *ADH1*, other alcohol dehydrogenase genes seem to have been horizontally acquired by W/S clade species (Figure 3-source data 1), including putative *ADH6* and *SFA1* orthologs, which also participate in alcoholic fermentation in *S. cerevisiae* (Drewke, Thielen, & Ciriacy, 1990; Ida, Furusawa, Hirasawa, & Shimizu, 2012). In all cases, except for *SFA1*, the “native” yeast orthologs appear to have been lost in W/S-clade genomes. This implies that ethanol production is conducted entirely by alcohol dehydrogenases of bacterial origin in W/S-clade species, which is, to our knowledge, a unique condition in fungi. We therefore wondered if more could be learned concerning the evolutionary history of these genes. To address this question, detailed phylogenetic analyses were conducted for Adh1 and Adh6, which are pivotal for alcoholic fermentation in yeasts. Maximum likelihood phylogenies were re-constructed using the top 4.000 phmmer hits obtained using *St. bombicola* Adh1 and Adh6 proteins as queries. The resulting Adh1 tree (Figure 3A) includes Adh1 protein sequences from both bacteria and fungi. Interestingly, all W/S-clade species clustered with strong support with the Acetobacteraceae (Proteobacteria) Adh1 proteins (Figure 3A and Figure 3B).

Within the Adh1 W/S-clade cluster, the overall phylogenetic relationships were in line with the expected relationships between the species (Figure 2-figure supplement 1), suggesting that a single HGT event occurred in the MRCA of this clade. This finding is also supported by the partial microsynteny conservation around the *ADH1* genes (Figure 3-figure supplement 1) among these species (*St. bombicola*, *C. magnoliae*, *St. bacillaris*, *C. apicola*, and *W. domercqiae*) and by the presence of other genes of bacterial origin in the vicinity of the *ADH1* gene in four of the species examined. Topology comparison tests (AU) strongly supported the Adh1 HGT event to the W/S clade (*P-value* = 0.008), which, together with the robustly supported branch that clusters the W/S-clade xenologs with bacteria, the synteny analysis, and the AI results, provide strong evidence for HGT.

Interestingly, Adh1 sequences from the outgroup *C. infanticola* also clustered with those of proteins of bacterial origin. However, the two Adh1 sequences from *C. infanticola* are not similar to those of the W/S clade (Figure 3C), as might be expected if a single HGT event were responsible for the acquisition of the *ADH1* gene in both lineages. These sequences are instead grouped, albeit with weak support, with Adh1 sequences from the distantly related Lactobacillales and Enterobacteriales (Figure 3C), implying that an independent HGT event may have occurred in the *C. infanticola* lineage. Nonetheless, topology tests did not strongly support an independent origin for W/S and *C. infanticola* Adh1 proteins (*P-value* = 0.073).

An extended Adh6 ML phylogeny (Figure 4A) was re-constructed with the top 10.000 phmmer hits to show that the W/S-clade sequences are indeed Adh6 orthologs. In this phylogeny, W/S-clade sequences clustered with strong support (>95%) with Proteobacteria Adh6 sequences, within a large Adh6 cluster that also encompasses known fungal Adh6 proteins. Interestingly, while Adh1 W/S-clade sequences grouped with those of the Acetobacteraceae (Figure 3A and Figure 3B), the Adh6 sequences are more closely related to those of other bacterial families, as highlighted in Figure 4B. The phmmer search failed to uncover Adh6 sequences in the *C. infanticola* proteome. Absence of an *ADH6* ortholog was further confirmed by a tBLASTx search against the *C. infanticola* genome using Adh6 sequences from both *S. cerevisiae* (KZV09178.1) and *St. bombicola* as queries. This result suggests that both *ADH1* and *ADH6* were lost in the MRCA of *C. infanticola* and the W/S clade. Interestingly, the *ADH6* xenologs were apparently duplicated several times within each W/S-clade species (Figure 4A, Figure 4B and Figure 3-source data 1), *Starmerella bacillaris* and *C. magnoliae* harboring the most paralogs (four in total).

**Figure 4.**
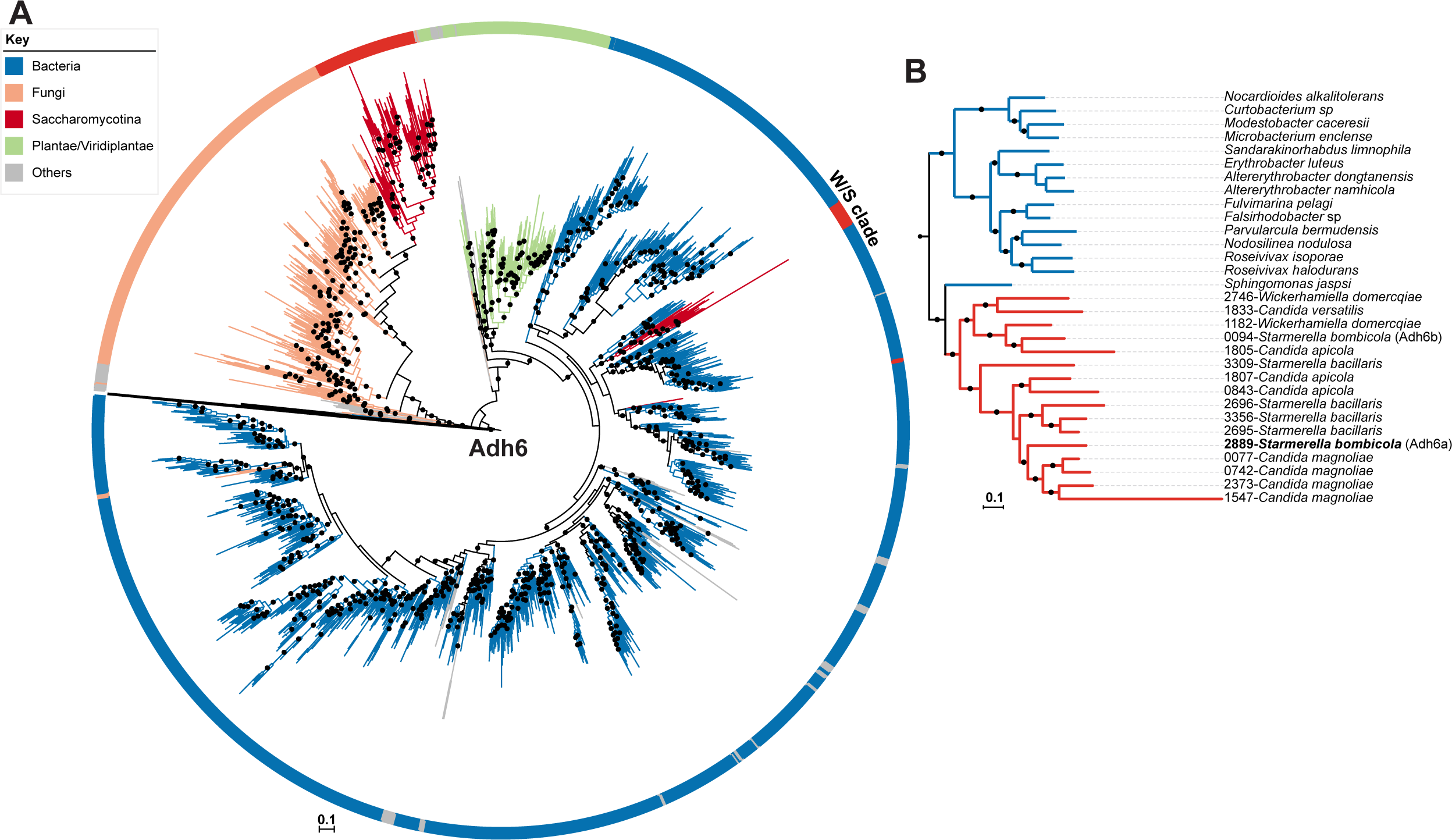
ML Phylogeny of Adh6 proteins. A) The phylogeny was constructed with the top 10.000 phmmer hits using *St. bombicola* Adh6 as a query (in bold, Panel B). Sequences with more than 80% similarity were eliminated. Branches with bootstrap support higher than 90% are indicated by black dots. Poorly represented lineages (<10 sequences) are shown in grey. Adh1-like sequences were collapsed as indicated. B) Pruned ML phylogeny depicting the phylogenetic relationship between Adh6 sequences from the W/S clade and their closest bacterial relatives.

### Functional role of alcohol dehydrogenase genes of bacterial origin in *St. bombicola*

All yeast (Saccharomycotina) species for which genome sequences are available harbor alcohol dehydrogenase orthologs, most notably *ADH1*, even if they do not produce ethanol (Gatter et al., 2016). This could suggest that acquisition of the bacterial alcohol dehydrogenase genes preceded the loss of the yeast ortholog, in which case the bacterial enzyme might be expected to be functionally different from its yeast counterpart, providing some new and advantageous characteristics that the yeast enzyme presumably lacked. The alternative hypothesis would be that *ADH* genes were first lost in the MRCA of *C. infanticola* and the W/S clade and were subsequently acquired from bacteria to restore alcoholic fermentation, possibly in connection to adaptation to a new environment. Hence, the elucidation of putative functional differences between the native yeast Adh1 and a bacterial xenolog would provide clues concerning possible evolutionary advantages of the “replacement” of the former by the latter. To investigate these potential differences, we characterized alcohol dehydrogenase (Adh) activity in three W/S-clade species, in the closely related yeast *B. adeninivorans* and in the model species *S. cerevisiae*, as well as in a distantly related fructophilic yeast species *Z. kombuchaensis*. All W/S-clade species tested were capable of using ethanol as carbon and energy source and have a Crabtree-negative behavior when growing on sugars, meaning that ethanol production in aerated batch cultures starts only when cell densities are very high, limiting oxygen availability (typically OD 30, ~5–10 g/L ethanol in *St. bombicola*).

In all non-W/S-clade species, the characteristic NADH-dependent Adh activity was readily observed, but no NADPH-dependent Adh activity was detected (Figure 5-figure supplement 1A), in line with available information concerning yeast enzymes (Cho & Jeffries, 1998; Dashko, Zhou, Compagno, & Piskur, 2014; Ganzhorn, Green, Hershey, Gould, & Plapp, 1987; Leskovac, Trivic, & Pericin, 2002). Conversely, all W/S-clade species tested (*St. bombicola*, C*. magnoliae*, and *St. bacillaris*) exhibited Adh activity when either NADH or NADPH was added to the reaction mixture (Figure 5-figure supplement 1B). When acetaldehyde was used as a substrate, the activity with NADH as cofactor was higher than with NADPH for all species tested. In fact, although both cofactors can be used for conversion of acetaldehyde into ethanol, there is a lower affinity for the substrate (higher K_m_) for NADPH-dependent activity in *St. bombicola* (Figure 5-figure supplement 1C). Interestingly, in *Acetobacter pasteurianus* a bacterial species in the Acetobacteraceae, the same family as the likely donor of W/S-clade Adh1, alcohol dehydrogenase activity was found to be NADH-dependent, although it is unclear whether NADPH was tested as a cofactor (Masud, Matsushita, & Theeragool, 2011).

**Figure 5.**
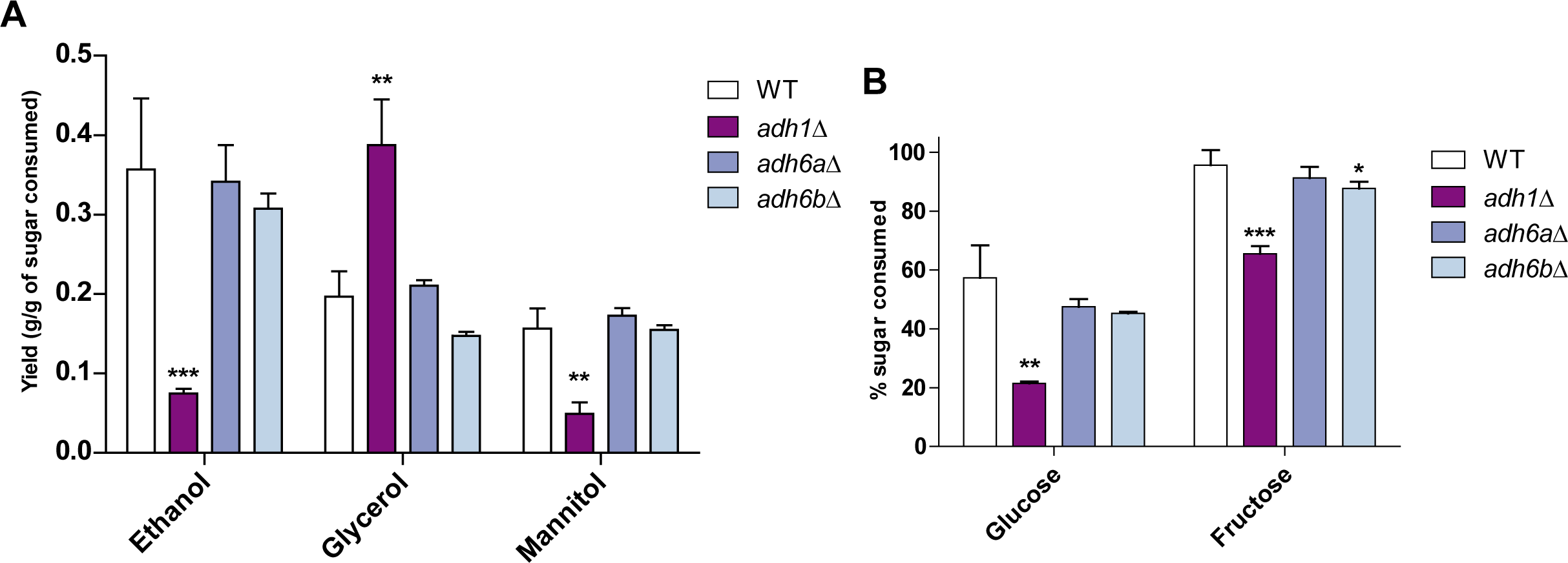
Metabolite production and sugar consumption in St. bombicola Wild Type (WT) and deletion mutants (adh1Δ, adh6aΔ, adh6bΔ). A) Ethanol, glycerol, and mannitol yields determined after 72 hours of growth. B) Percentage of sugar (fructose and glucose) consumed after 72 hours of growth. Error bars represent standard deviation of assays performed in duplicate in two biological replicates. All strains were grown in YP supplemented with 10% (w/v) fructose and 10 % (w/v) glucose at 30ºC with aeration. Statistically significant differences (student’s t-test, two-tailed) between WT and deletion mutants for sugar consumption and metabolite production are shown (** P-value < 0.01; *** P-value < 0.005).

In *S. cerevisiae*, the paralogous enzymes Adh1, Adh2, Adh3, and Adh5 are mainly responsible for the inter-conversion of ethanol and acetaldehyde in a NADH-dependent manner. Adh6 and Adh7 are also capable of catalyzing this reaction, but they use NADPH instead (de Smidt et al., 2008). Since both *ADH1-* and *ADH6-*like genes are present in the genomes of all W/S-clade species studied, it was not clear which enzyme (Adh1-type or Adh6-type) was responsible for the NADPH-dependent inter-conversion of ethanol and acetaldehyde observed in W/S-clade species. To elucidate this, and to evaluate the impact of alcohol dehydrogenases on metabolism, three knock-out mutants were constructed in *St. bombicola (adh1∆*, *adh6a∆*, and *adh6b∆*). During aerated growth, deletion of *ADH1 (adh1∆*) did not seem to significantly affect specific growth rates in glucose or fructose when compared to the WT (Figure 5-figure supplement 2A). However, we noted a five-fold decrease in ethanol production (Figure 5A) and the absence of growth on ethanol as sole carbon and energy source in the *adh1∆* mutant. Although some ethanol was produced, no Adh activity was detected in cell-free extracts of the *adh1∆* mutant when either NADH or NADPH was used. These results suggest that Adh1 is the main enzyme used in alcoholic fermentation in *St. bombicola* and that it therefore likely accepts both NADH and NADPH as cofactors. Interestingly, the significant decrease in ethanol yield seems to be counterbalanced by a concomitant increase in glycerol production (Figure 5A), similarly to what has been observed in the *S. cerevisiae adh1∆* mutant (de Smidt, du Preez, & Albertyn, 2012). Moreover, growth of the *adh1∆* mutant cultivated on identical growth medium but under limited aeration was severely affected (Figure 5-figure supplement 2B). Taken together, the two observations strongly suggest that Adh1 plays an important role in redox homeostasis, namely in NAD^+^ regeneration in the absence of oxygen, because, similarly to *S. cerevisiae*, glycerol formation in *St. bombicola* is probably a NAD^+^ regenerating reaction. Consistent with this hypothesis, we did not detect NADP^+^ dependent glycerol dehydrogenase activity in cell-free extracts (Figure 5-figure supplement 1D), and in at least one W/S-clade species, this reaction was shown to be NADH-dependent (Van Bogaert, De Maeseneire, Develter, Soetaert, & Vandamme, 2008). In contrast, mannitol production, which is a NADP^+^ regenerating reaction (Lee, Koo, Kim, & Hyun, 2003), was significantly decreased in the *adh1∆* mutant (Figure 5A). The deletion of each of the two *ADH6* paralogous genes *(adh6a∆* and *adh6b∆* mutants) did not significantly affect ethanol production (Figure 5A) or consumption. This means that, although some of the enzymes encoded by these genes might be involved in the production of ethanol in the absence of Adh1, when Adh1 is functional, they are not essential for ethanol metabolism and are probably mainly involved in other metabolic reactions. Finally, to ascertain how perturbations in alcoholic fermentation might affect the relative preference of *St. bombicola* for fructose over glucose, we monitored the consumption of both sugars in aerated cultures over time in the *adh1∆, adh6a∆* and *adh6b∆* mutants. There was a significant decrease in sugar consumption rates in the *adh1∆* mutant (Figure 5B), but fructophily was still observed in all mutants, which leads to the conclusion that the xenolog alcohol dehydrogenases do not affect sugar preference in this species.

### Horizontally transferred *SUC2* is essential for sucrose assimilation in *St. bombicola*

*SUC2* encodes the enzyme invertase and is an essential gene for sucrose assimilation in yeasts (Carlson et al., 1981). Invertase (Suc2) extracellularly hydrolyzes this disaccharide into fructose and glucose. In *S. cerevisiae*, the *MAL* and the *IMA* genes have also been shown to play a role in sucrose hydrolysis (Deng et al., 2014; Voordeckers et al., 2012), but these genes are absent in the genomes of W/S-clade species, meaning that the horizontally transferred invertase appeared to be the only enzyme with sucrose-hydrolyzing capacity encoded in the W/S-clade genomes investigated. In a phylogenetic tree reconstructed using the 200 top phmmer hits to 2301–5490- *Candida magnoliae* (one of the W/S-clade species investigated), strong support for clustering of W/S-clade sequences to Acetobacteraceae SacC sequences was observed (Figure 6A and Figure 6B). A topology comparison test (AU) also strongly supported the HGT hypothesis (*p-value* = 4e-07).

The phylogeny shown in Figure 6A is in line with SacC being the Suc2 homolog in bacteria, which is also corroborated by the fact that it is known to hydrolyze sucrose, as well as other β-fructans (Martel et al., 2011; Wanker, Huber, & Schwab, 1995). Notably, the same group of Acetobacteraceae species clustered with W/S-clade species in the Adh1 phylogeny (Figure 3B and Figure 6B), suggesting that both genes may have been acquired in a single HGT event (Figure 7). We also noted that the closest relative of the W/S clade, *C. infanticola*, lacks a *SUC2* gene.

**Figure 7.**
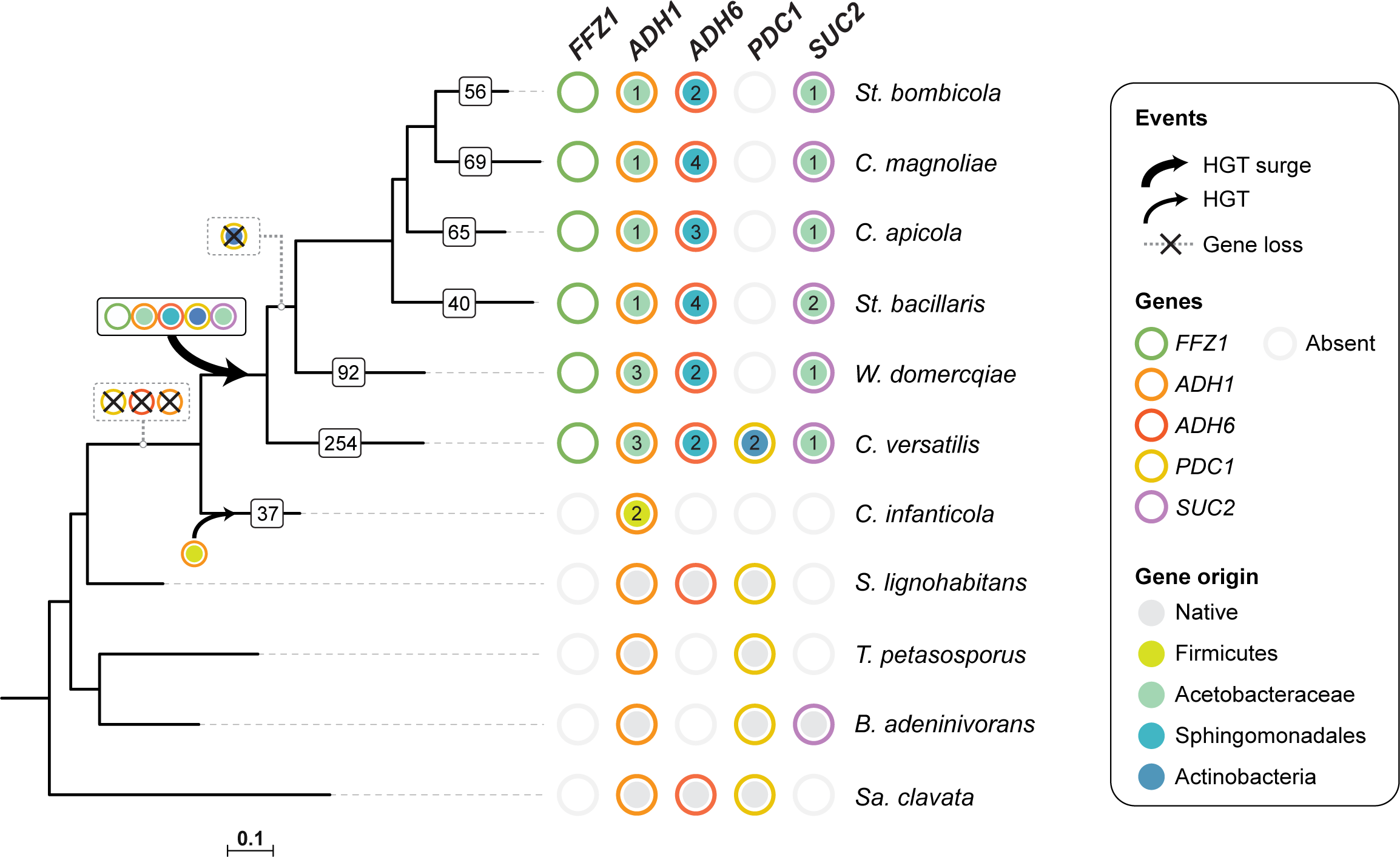
Loss and acquisition of sugar metabolism related genes in the W/S clade and closely related lineages. The number of genes of bacterial origin (AI>0.1) are indicated for each W/S-clade branch (white squares) and for the closest relative C. infanticola. For each of four genes, *ADH1*, *ADH6*, *SUC2*, and *PDC1*, presence, absence, and the native or bacterial origin of the orthologs found in the cognate draft genomes are shown for each species next to the respective branch of the tree. Each gene is represented by circles with different line colors (green for previously studied *FFZ1* (Goncalves et al., 2016), orange for *ADH1*, red for *ADH6*, yellow for *PDC1* and purple for *SUC2*). For xenologs, the different predicted bacterial donor lineages are denoted by different fill colors as indicated in the key. For W/S-clade species, the number of paralogs found in the cognate draft genome is also shown. Inferred gene losses (cross) and HGTs events (arrows) are indicated in the tree using the same notation for each orthologous group.

To ascertain whether the horizontally acquired invertase gene is responsible for sucrose assimilation in the W/S clade, a *sacC* deletion mutant (referred henceforth to as *suc2∆*) was constructed in *St. bombicola*. Growth assays in medium supplemented with sucrose as sole carbon and energy source, showed that the two *suc2∆* strains were unable to grow (Figure 6- figure supplement 1A), while the WT strain attained high cell densities. Furthermore, the *suc2∆* mutant failed to consume measurable amounts of sucrose, even after 72 hours (Figure 6-figure supplement 1A). Interestingly, when the WT strain grew on sucrose, the decrease in sucrose concentrations was accompanied by the appearance of fructose and glucose in the growth medium, strongly suggesting that the horizontally transferred *SUC2* gene encodes an extracellular invertase (Figure 6-figure supplement 1B). This conclusion is consistent with the apparent absence of genes encoding sucrose transporters in the W/S-clade genomes analyzed, which indicates that sucrose must be first hydrolyzed outside the cell to be used as a carbon and energy source.

## Discussion

The early-derived yeast lineage, here named the *Wickerhamiella/Starmerella* (W/S) clade, comprises several species that have previously attracted attention due to their unusual metabolic features. The most prominent example is *St. bombicola*, a species used for the production of sophorolipids, which are amphipathic molecules that are employed as biosurfactants (Samad, Zhang, Chen, & Liang, 2015; Takahashi et al., 2011). *Starmerella bacillaris* is often found in wine fermentations and is known for diverting an important fraction of its carbon flux towards the production of glycerol instead of ethanol (Englezos et al., 2015). *Candida magnoliae* has been reported to be capable of producing large amounts of erythritol (Ryu, Park, Park, Kim, & Seo, 2000). More recently, we reported that fructophily was an important common trait that unified these species and all others belonging to the W/S clade, and we also consubstantiated a strict correlation between the presence of the transporter Ffz1 and fructophily. Here we show that presence of the Ffz1 transporter is a pre-requisite for fructophily in *St. bombicola*, as previously observed in the *Z. rouxii* (Leandro et al., 2014). The stronger emphasis on the production of sugar alcohols as byproducts of metabolism at the expense of ethanol seemed to be also a common trait between the species examined (Lee, Song, & Kim, 2003; Ryu et al., 2000) and led us to hypothesize that the preference of these yeasts for fructose was likely to be part of a broader remodeling of metabolism connected to the adaptation to the high sugar environments in which these yeasts thrive. The present work reflects our effort to uncover other aspects of this adaptation using comparative genomics as a starting point.

Our examination of genes acquired from bacteria showed that the number of HGT events from bacteria into the W/S clade and its neighbor lineage (represented by the species *C. infanticola*), far exceeded the number of events reported for other Saccharomycotina lineages (Marcet-Houben & Gabaldon, 2010). The largest number of HGT events was detected in the earliest-derived species in the W/S clade, *C. versatilis*, which together with the phylogenetic signal in the genes that were acquired through HGT, suggests that a large surge of acquisitions probably occurred in the MRCA of the clade (Figure 7). Under this model, most extant lineages subsequently discarded a larger portion of the xenologs originally present in the common ancestor. We noted that the set of xenologs present in at least two extant W/S-clade species is enriched for genes encoding proteins that affect redox balance in the cell. In fact, changes in fluxes through main metabolic pathways were previously shown to impact redox balance and oxidative stress in yeasts (González-Siso, García-Leiro, Tarrío, & Cerdán, 2009), which is consistent with our hypothesis that associates the acquisition of bacterial genes with adaptive changes in metabolism. It seems possible that, in addition to HGT events common only to W/S-clade species, additional HGT events also took place in the MRCA of *C. infanticola* and the W/S clade because at least five from the 37 genes of apparent bacterial origin in *C. infanticola* also have bacterial origin in the W/S clade. While the bacterial donor lineage seems to be quite different between *C. infanticola* and the W/S clade for one gene (*ADH1*), strongly suggesting that they were acquired in separate events, the other four genes seem to have originated from the same donor lineage, possibly pointing to a single event. The remaining HGT-derived genes (32) seem to be specific to *C. infanticola*.

The most striking finding concerning the function of the transferred genes is the profound remodeling of the ubiquitous alcoholic fermentation pathway, which is used by yeasts to convert pyruvate into ethanol with concomitant regeneration of NAD^+^. The first step of the pathway, the conversion of pyruvate into acetaldehyde, is normally catalyzed by the enzyme Pdc1, but in most W/S-clade yeasts, it is apparently conducted solely by a related decarboxylase encoded in *S. cerevisiae* by the *ARO10* gene. In *S. cerevisiae*, phenylpyruvate is the primary substrate of Aro10, which links this enzyme to amino acid metabolism, rather than alcoholic fermentation (Romagnoli, Luttik, Kotter, Pronk, & Daran, 2012; Vuralhan et al., 2005). Interestingly, the earliest-derived W/S-clade species among those examined, *C. versatilis*, also lacks a yeast-like *PDC1* gene but possesses two genes of bacterial origin encoding *PDC1* orthologs, which coexist with *ARO10.* In this species, it is possible that the xenologs, and not *ARO10*, carry out the conversion of pyruvate into acetaldehyde.

The second step in alcoholic fermentation is the conversion of acetaldehyde into ethanol, which is conducted in *S. cerevisiae* mainly by Adh1. Again, “yeast-like” *ADH1* genes were absent from all W/S-clade genomes examined, and it seems that the MRCA of the W/S clade acquired a bacterial *ADH1* gene, from which extant *ADH1* xenologs found in all extant W/S-clade species examined were derived. A similar occurrence was detected involving the loss of the “yeast-like” *ADH6* ortholog, encoding a NADPH-dependent branched chain alcohol dehydrogenase and the acquisition of bacterial orthologs, although the bacterial donor lineages of the *ADH1* and *ADH6* xenologs seem to be distinct. We showed that, in *St. bombicola*, the *ADH1* xenolog is absolutely required for growth on ethanol and is also mainly responsible for alcoholic fermentation, thereby performing the functions typically fulfilled by two different enzymes in *S. cerevisiae* (where *ADH2* catalyzes ethanol assimilation). The two *ADH6* xenologs played a minor role, if any, in alcoholic fermentation. Interestingly, fructophilic bacteria have also undergone a remodeling of their alcohol dehydrogenases, but as far as we could ascertain, in the fructophilic yeast *St. bombicola*, there was no link between fructophily and the presence of *ADH* xenologs, since fructophily was unaffected in the alcohol dehydrogenase mutants. It remains to be investigated if the preference for fructose over glucose in yeasts is linked to a possible role of fructose as an electron acceptor, like in fructophilic bacteria.

The events leading to the observed profound remodeling of the alcoholic fermentation pathway in W/S-clade yeasts are impossible to retrace with certainty because of their antiquity. However, we may outline at least two distinct plausible hypotheses. One possibility might be that the genes required for alcoholic fermentation, *ADH1* and *PDC1*, as well as *ADH6*, were lost in an ancestral event, probably in the MRCA of the W/S clade and *C. infanticola*, possibly because cofactor recycling through alcoholic fermentation became superfluous. Since all yeast species examined so far, except those included in the W/S clade, possess “yeast-like” *ADH1* and *PDC1* orthologs, this occurrence can be considered a very unusual, if not unique event, in the Saccharomycotina. Later, presumably as part of the adaptation to the floral niche, bacterial *ADH* and *PDC1* genes were acquired from bacteria, restoring alcoholic fermentation. The ecological opportunity for this horizontal exchange of genetic material seems to have existed for a long time, since W/S clade Adh1 proteins are most closely related to those of a bacterial lineage that is also frequently associated to the floral niche (Acetobacteraceae) (Iino et al., 2012; Suzuki et al., 2010; Tucker & Fukami, 2014). While *ADH1* and *ADH6* xenologs persist in all extant W/S-clade species examined, bacterial *PDC1* xenologs, if they were indeed also acquired by the MRCA of the clade, were subsequently lost in most species. These hypothesized losses may have occurred as Aro10 presumably evolved to fulfill the function of Pdc1. *C. versatilis*, which is the earliest-derived species in the W/S clade, is a notable exception that possesses two *PDC1* xenologs in addition to *ARO10*. One argument against the loss scenario might be that no extant yeast species has yet been found devoid of at least one *ADH1* ortholog, irrespective of its origin. In fact, even the model yeast *Y. lipolytica*, which does not produce ethanol, possesses alcohol dehydrogenases of the *ADH1*-type that are involved in ethanol assimilation (Gatter et al., 2016). Therefore, we must assume that,if a yeast lineage devoid of *ADH1* ever existed, it has either become extinct or it has not been sampled yet.

An alternative scenario would be that the bacterial genes were acquired by lineages that still had the “yeast-like” versions of the *ADH1*, *ADH6*, and *PDC1* genes because they brought along new adaptive traits. This scenario, in which the “yeast-like” versions of the genes would have been lost after acquisition of the bacterial version, does not call for the existence, extant or in the past, of yeast lineages devoid of alcohol dehydrogenases. However, it relies on the ability of the bacterial genes to improve fitness of the recipient lineage, presumably the MRCA of the W/S clade, possibly as it adjusted to a new environment. Our assessment of the performance of the Adh1 enzymes in W/S-clade species showed that they provide a potentially advantageous characteristic when compared to their yeast counterparts, since they are capable of regenerating both NAD^+^ and NADP^+^, while the yeast enzymes accept only NADH as a cofactor. However, at least in *St. bombicola*, elimination of Adh1 was compensated by an increase in glycerol formation, which is a NAD^+^ regenerating reaction, and not of mannitol formation, which regenerates NADP^+^. These results suggest that the role of Adh1 in *St. bombicola* is mainly related to NAD^+^/NADH homeostasis, as has been observed in the corresponding *S. cerevisiae* mutant. This interpretation is consistent with the lower K_m_ for the substrate observed when NADH was used as a cofactor. However, it is possible that this observation in *St. bombicola* does not reflect the situation in the ancient recipient lineage in which the acquisition of the bacterial genes took place, and that the ability to regenerate both oxidized cofactors was indeed pivotal in the process, which may also hold true for the scenario in which loss preceded acquisition of the bacterial xenologs. Therefore, the presence of xenologs involved in alcoholic fermentation in extant W/S-clade yeasts can be explained by the need to restore alcoholic fermentation in a lineage that had previously lost it, in which case the involvement of bacterial genes may have been circumstantial and a consequence of the availability of the donor in the same environment. Loss and subsequent reacquisition of a metabolic pathway through multiple HGT events was reported for the unicellular red algae *Galdieria phlegrea* where massive gene loss occurred concomitantly with adaptation to a specialized niche in the common ancestor of Cyanidiophytina red algae (Qiu et al., 2013). Taking all data into account, we posit that a similar scenario in which *ADH* and *PDC1* gene loss precedes acquisition (Figure 7) is most consistent with the evidence we gathered. However, it cannot be excluded that bacterial genes may have been selected because they encode enzymes that possess distinct catalytic properties when compared with their yeast counterparts, although we did not find direct evidence for that in the extant species examined.

*SUC2* is among the genes acquired specifically by the MRCA of the W/S clade. Unlike the “yeast-like” version of *ADH1*, which is ubiquitous in the Saccharomycotina, except in the W/S clade, evolution of “yeast-like” *SUC2* seems to be punctuated by multiple loss events resulting in a patchy extant distribution (Carlson, Celenza, & Eng, 1985), which is probably linked to its role as a “social” gene (Sanchez & Gore, 2013). The acquisition of the invertase gene from bacteria (the same donor lineage as *ADH1*) is consistent with the idea that a major remodeling of sugar metabolism took place in the MRCA of the W/S clade, as it adapted to the sugar-rich floral niche. Intriguingly, the *SUC2* phylogeny suggests that fungal invertases may be derived from a common ancestor, itself of bacterial origin (Figure 6A).

In summary, our results uncovered to our knowledge for the first time, an instance of major remodeling of central carbon metabolism in fungi involving the acquisition of a multitude of bacterial genes, among which an alcohol dehydrogenase with novel catalytic properties. Taken together with previous observations concerning unusually high production of sugar alcohols and/or lipids by W/S-clade species, our results suggest that the underpinnings of central metabolism are very different in these yeasts when compared with the model species *S. cerevisiae*, in particular in what concerns the mechanisms of redox homeostasis in which fructose may participate as electron acceptor, as observed for fructophilic bacteria.

## Materials and Methods

### Strains

Yeast strains were obtained from PYCC (Portuguese Yeast Culture Collection, Caparica, Portugal). All strains were maintained in YPD medium.

### Comparative genomics of sugar metabolism related genes

A custom query consisting of *S. cerevisiae* genes related to sugar metabolism (pentose phosphate pathway, glycolysis, and ethanol fermentation) was constructed. To retrieve central carbon metabolism genes in fructophilic yeasts, a tBLASTx search was performed against W/S-clade genomes (*C. versatilis* JCM 5958, *St. bombicola* PYCC 5882, *St. bacillaris* 3044, *C. magnoliae* PYCC 2903, *C. apicola* NRRL Y-50540 and *W. domercqiae* PYCC 3067), and *Zygosaccharomyces kombuchaensis* CBS 8849 using this query. The genomes of *Sugiyamaella lignohabitans* CBS 10342, and *Blastobotrys adeninivorans* LS3, which are the most closely related species to the W/S clade were added to the database. The best tBLASTx hit sequences retrieved from the analysis were subsequently blasted against the NCBI non-redundant protein database, and orthology was assumed whenever the best hit in *S. cerevisiae* was the corresponding protein in the query.

Genomes of *C. apicola* (Vega-Alvarado et al., 2015) and *C. versatilis* (PRJDB3712) are publicly available, while the draft genomes of the remaining W/S-clade species examined and of *Z. kombuchaensis* were generated in the course of a previous study (Goncalves et al., 2016) and are also publicly available (Bioprojects PRJNA 416500 and PRJNA 416493 or https://figshare.com/s/905b12b0ac9bf7c46b8f).

For species included in Figure 7 (*Trichomonascus petasosporus* and *Saprochaete clavata*), the presence or absence of sugar metabolism related genes *(ADH1*, *ADH6*, *PDC1* and *SUC2*) were assumed depending on whether the best BLASTp and tBLASTx hits (E-value <e-10) against the respective genome databases was the respective ortholog in *S. cerevisiae*.

### Protein prediction in newly sequenced genomes

For *St. bombicola* PYCC 5882, *St. bacillaris* PYCC 3044, *C. magnoliae* PYCC 2903, *W. domercqiae* PYCC 3067, and *C. infanticola* DS-02 (PRJNA318722) the complete proteome was predicted with AUGUSTUS (Stanke, Diekhans, Baertsch, & Haussler, 2008) using the complete model and *S. cerevisiae*, *Scheffersomyces stipitis*, and *Y. lipolytica* as references. To assess the completeness of the predicted proteomes in each case, the number of predicted proteins was compared to the number of proteins reported for the annotated proteomes of *W. domercqiae* JCM 9478 (PRJDB3620) and *S. bombicola* JCM 9596 (PRJDB3622). Predictions that used *S. cerevisiae* as a reference turned out to be the most complete (~4.000 predicted proteins) and were therefore used for all downstream analyses. For *C. apicola* NRLL Y-50540 and *C. versatilis* JCM 5958, the publicly available proteome was used.

### High throughput detection of HGT-derived genes from bacteria in the W/S clade

Query protein sequences were searched against a local copy of the NCBI refseq protein database (downloaded May 5, 2017) using phmmer, a member of the HMMER3 software suite (Eddy, 2009) using acceleration parameters --F1 1e-5 --F2 1e-7 --F3 1e-10. A custom perl script sorted the phmmer results based on the normalized bitscore (*nbs*), where *nbs* was calculated as the bitscore of the single best-scoring domain in the hit sequence divided by the best bitscore possible for the query sequence (i.e., the bitscore of the query aligned to itself). The top ≤ 10,000 hits were retained for further analysis, saving no more than five sequences per unique NCBI Taxonomy ID.

The alien index score (AI) was calculated for each query protein (modified from Gladyshev et al. 2008)(Gladyshev et al., 2008). Two taxonomic lineages were first specified: the RECIPIENT into which possible HGT events may have occurred (Saccharomycetales, NCBI Taxonomy ID 4892), and a larger ancestral GROUP of related taxa (Fungi, NCBI Taxonomy ID 4751). The AI is given by the formula: AI = (*nbsO* − *nbsG*), where *nbsO* is the normalized bitscore of the best hit to a species outside of the GROUP lineage, *nbsG* is the normalized bitscore of the best hit to a species within the GROUP lineage (skipping all hits to the RECIPIENT lineage). AI can range from 1 to −1. AI is greater than zero if the gene has a better hit to a species outside of the group lineage and can be suggestive of either HGT or contamination.

Phylogenetic trees were constructed for all HGT candidates with AI > 0.1. Full-length proteins corresponding to the top 200 hits (E-value < 1 × 10^−10^) to each query sequence were extracted from the local database using esl-sfetch (Eddy, 2009). Sequences were aligned with MAFFT v7.310 using the E-INS-i strategy and the BLOSUM30 amino acid scoring matrix (Katoh & Standley, 2013) and trimmed with trimAL v1.4.rev15 using its gappyout strategy (Capella-Gutierrez, Silla-Martinez, & Gabaldon, 2009). Genes with trimmed alignments < 150 amino acids in length were excluded. The topologies of the remaining genes were inferred using maximum likelihood as implemented in IQ-TREE v1.5.4 (Nguyen, Schmidt, Haeseler, & Minh, 2015) using an empirically determined substitution model and rapid bootstrapping (1000 replications). The phylogenies were midpoint rooted and branches with local support less than 95 were collapsed using the ape and phangorn R packages (Paradis, Claude, & Strimmer, 2004; Schliep, 2011). Phylogenies were visualized using ITOL version 3.0 (Letunic & Bork, 2016).

From the total of 625 phylogenies, the best HGT candidates were selected according to two criteria: the putative bacterial ortholog should be present in, at least, two W/S-clade species and they should cluster with a bacterial group with strong bootstrap support (>90%). Putative HGT-derived genes shared with non-W/S-clade species were excluded. Application of these criteria selected 200 strong candidate trees (Figure 3- source data 1), many of which referred to the same ortholog (in cases where the gene is present in more than one species). A final number of 52 different ortholog groups was established after collapsing the replicate phylogenies and lineage-specific gene duplications. The resulting set of 52 orthologs was subsequently cross-referenced with GO and InterPro annotations provided by InterproScan and the Joint Genome Institute MycoCosm Portal (Grigoriev et al., 2014). KEGG annotation was performed using the KAAS database (Moriya, Itoh, Okuda, Yoshizawa, & Kanehisa, 2007). A BLAST KOALA (Kanehisa et al., 2016) was also conducted in the final dataset (52 proteins).

### Phylogenetic analyses of Pdc1, Adh1, Adh6, Suc2 and topology tests

The top 500 NCBI BLASTp hits from searches against the non-redundant database using Pdc1 from *S. cerevisiae* (CAA97573.1)*, St. bombicola* Pdc1-like, and *C. versatilis* Pdc1-like from apparent bacterial origin as queries, were selected. Sequences with more than 90% similarity were removed using CD-HIT v 4.6.7 (Li & Godzik, 2006). Pdc1 sequences from the closest relative species *Trichomonascus petasosporus* (JGI), *Blastobotrys adeninivorans* LS3 (JGI) and *Saprochaete clavata* CNRMA 12.647 (Genbank) of the W/S clade were also added to the alignment. A total of 479 proteins were aligned using MAFFT v 7.2.15, (Katoh & Standley, 2014) using the fast but progressive method (FFT-NS-2). Poorly aligned regions were removed with trimAl (Capella-Gutierrez et al., 2009) using the “gappyout” option. The ML phylogeny in Figure 2 was constructed with IQ-TREE v 1.4.3 (Nguyen et al., 2015) using the LG+I+G substitution model. For the Suc2 phylogeny, the top 200 hits from the phmmer search against the local database were selected and the ML phylogeny was constrcted as above. Given the absence of other Saccharomycotina sequences in the top 200 hits for Adh1, the top 4000 hit sequences were selected instead. For Adh6, no Saccharomycotina sequences were found even in the top 4000 phmmer hits, so the top 10.000 top hit sequences were used in this case. For both phylogenies, CD-HIT (Li & Godzik, 2006) was used to remove sequences with more than 85% (Adh1) and 80% (Adh6) similarity. For Adh1, a preliminary ML tree was constructed to eliminate sequences outside the Adh1 family. A final set of 976 Adh1 sequences were subsequently aligned and trimmed as aforementioned and used to construct the final Adh1 phylogeny. ML phylogenies were constructed as previously. For all phylogenies, the different lineages are highlighted with distinct colors: red for Saccharomycotina, blue for Bacteria (which includes several different lineages) and orange for Fungi (which represents fungal lineages other than Saccharomycotina). Original raw phylogeny files can be accessed using the following links: https://figshare.com/s/c59f135885f31565a864. The likelihood of HGT for *ADH1* and *SUC2* was investigated assuming monophyly of Saccharomycotina as the constrained topology (Figure 3-figure supplement 2A and Figure 3-figure supplement 2C). The best tree was inferred in RAxML, and ML values for constrained and unconstrained trees were also calculated. The AU test (Shimodaira, 2002) implemented in CONSEL (Shimodaira & Hasegawa, 2001) was used to compare the unconstrained best tree and the best tree given a constrained topology. To test the independence of Adh1 acquisition, monophily of W/S clade and *C. infanticola* was assumed (Figure 3-figure supplement 2B). Phylogenies were visualized using iTOL version 3.0 (Letunic & Bork, 2016).

### Detection of enzymatic activities in cell-free extracts

Cultures were grown overnight in YPD medium until late exponential phase (OD_640nm_ ~15–25). Cells were then collected by centrifugation (3,000 x g for 5 minutes), washed twice with cold Tris-HCL buffer (pH=7.6), and disrupted with glass beads in 500 µL of Lysis Buffer (0.1 M triethanolamine hydrochloride, 2 mM MgCl_2_, 1 mM DTT and 1 µM PMSF) with six cycles of 60 seconds vortex-ice. Cell debris were removed by centrifugation at 4ºC, 16000 x g for 20 minutes and the extracts were stored at −20ºC. Alcohol dehydrogenase activity (Adh) assays were performed at 25ºC in 500 µL reaction mixtures containing 50 mM Potassium Phosphate buffer (pH=7.5), 1 mM of NADH or NADPH, and 25 µL of cell-free extract. The reaction was started by adding acetaldehyde to a final concentration of 100 mM, and reduction of NADH and NADPH was monitored spectrophotometrically by the decrease in absorbance at 340 nm for two minutes. For *St. bombicola* PYCC 5882, 5 mM, 12.5 mM, 50 mM, and 100 mM as final concentrations of acetaldehyde were also used. The absence of Adh activity in the *adh1∆* mutant was also confirmed with protein extract of *adh1∆* cells grown in 20FG medium (conditions where ethanol was detected by HPLC in the mutant, Figure 5A), using up to three times more protein extract.

For the detection of NADP^+^ dependent glycerol dehydrogenase activity (Klein, Swinnen, Thevelein, & Nevoigt, 2017) in *St. bombicola*, cultures and cell-free extracts were obtained as above. Glycerol dehydrogenase activity was measured in a reaction mixture containing 50 mM Tris-HCl (pH=8.5) buffer, 1 mm NADP^+^ and 25 µL of cell-free extract. The reaction was started by adding glycerol to a final concentration of 100 mM of, and NADPH formation was monitored spectrophotometrically for two minutes.

### Construction of *St. bombicola* deletion mutants

Standard molecular biology techniques were performed essentially as described in (Sambrook & Russell, 2001) using *E.*coli DH5α as host. *St. bombicola* PYCC was used in all procedures involving this species. Disruption constructs were designed essentially as outlined by Van Bogaert et al (2008)(Van Bogaert et al., 2008). The *St. bombicola GPD* promoter was first amplified by PCR and fused to the hygromycin B phosphotransferase gene (*hygB*) from *E.coli* and the *CYC1* terminator from *S. cerevisiae*. Phusion High Fidelity (Thermo Fisher Scientific, Waltham, Massachusetts, USA) was used for *hygB* and *GPD* promoter amplifications. A 491-bp fragment of the GPD promoter (Van Bogaert et al., 2008) was amplified using the primer pair GPD_SacI_Fw/GPD_Hind III_Rv (Figure 5-source data 2). The *TEF1* promoter from p414TEF-CYC (Mumberg, Muller, & Funk, 1995) vector was replaced by the *GPD* promoter using *Sac* I and *Hin*d III. The primer pair Hyg_Hind III_Fw/Hyg_Xho I_Rv was used to amplify the *hygB* gene from a commercial plasmid (pBlueScript-hyg, (Niklitschek et al., 2008)). The amplicon was then cloned into the previously obtained p416GPD-CYC plasmid. The resulting plasmid harbors the hygromycin resistance gene controlled by the *St. bombicola GPD* promoter and followed by the *CYC1* terminator of *S. cerevisiae*.

For disruption of the *ADH1*, and *SUC2* genes the coding sequences (CDS) with 1kb upstream and downstream were amplified from genomic DNA using the primer pairs listed in (Figure 5-source data 2). The two fragments, corresponding to each of the genes, were separately cloned into the PJET1.2 plasmid. The GPD-HYG-CYC cassette was then cloned into each of the resulting PJET1.2 plasmids using the restriction enzymes listed in Figure 5-source data 2. For *FFZ1*, *ADH6*a, and *ADH6b*, two sets of primers were used to amplify 1 kb upstream and 1 kb downstream of the CDS. The upstream and downstream fragments of each of the genes were subsequently cloned into the p416GPD-HYG-CYC plasmid using suitable enzymes (Figure 5-source data 2), yielding three plasmids each containing a distinct disruption cassette.

*St. bombicola* was transformed by electroporation with each gene disruption construct in turn, amplified by PCR from the respective plasmid template using Phusion High Fidelity DNA polymerase and the primers listed in (Figure 5-source data 2), using the protocol described by Saerens KM et al., 2011 (Saerens, Saey, & Soetaert, 2011). Two different transformants from each gene disruption transformation were subsequently used for all phenotypic assays.

### Sugar assimilation, consumption, and metabolite production assessments

For metabolite and sugar consumption profiling, 10-mL cultures of *St. bombicola* (WT and mutants) were grown overnight at 30ºC with orbital shacking (180 rpm) in YP medium supplemented with different concentrations of fructose and glucose. The overnight culture was used to inoculate a fresh culture in 30 mL of the same medium to an OD_640nm_ of 0.2, which was incubated under the same conditions. Growth was monitored until late stationary phase was reached (typically after 150 hours), and 2-mL samples were taken at several time points, centrifuged at 12000g for 5 min, and analyzed by HPLC, as previously described (Goncalves et al., 2016). Statistical significance was tested using a two-tailed student´s t-test, implemented in GraphPad Prism v5. Growth on sucrose and on ethanol was assessed in wild type and mutants (*suc2∆*, *adh1∆*, *adh6a∆*, and *adh6b∆*) cultivated overnight in YP medium supplemented with 2% (w/v) sucrose (*suc2∆*) or 2% (v/v) ethanol (*adh1∆*, *adh6a∆*, and *adh6b∆*) with orbital shacking (180 rpm) at 30ºC. Cultures were transferred into the same medium (OD_640nm_ = 0.2), and growth was monitored over time. Consumption of sucrose at different time points was monitored by HPLC.

### Growth assays

*St. bombicola* WT and *adh1∆* mutants were tested for growth aerobically (favoring respiration) and microaerophily (favoring fermentation) conditions. For the mutants, two biological replicates from independent transformations were used. For microaerophily experiments, a 24 hour pre-culture in SC medium supplemented with 0.2% glucose was performed. These cultures were used to inoculate 200 µL of SC medium supplemented with the desired carbon source (2% (w/v) glucose or 2% (w/v) fructose, at a 1:40 ratio in a 96 well plate. The absorbance of each well was read by an unshaken BMG FLUOstar Omega plate reader (Kuang, Hutchins, Russell, Coon, & Hittinger, 2016) every 120 min at 600 nm for five days. For aerobic growth, a 10 mL pre-culture in SC medium supplemented with 0.2% glucose was performed overnight with shaking (200 rpm). Cells were transferred to 30 mL (in a 250-mL flask) of SC medium supplemented with 2% (w/v) glucose or 2% (w/v) fructose until a final OD_600nm_=0.2. Cultures were grown for five days, with shaking.

---

**Figure 6-ML phylogeny of fungal Suc2 and bacterial SacC proteins.** A) ML phylogeny of Suc2/SacC proteins (top 200 phmmer hits). W/S-clade species are highlighted. Only branches with bootstrap support higher than 90% are indicated (black dots). B) Pruned ML phylogeny of Suc2 depicting the phylogenetic relationship between the W/S clade and Acetobacteraceae.

## Figure Supplements

**Figure 2-figure supplement 1.**
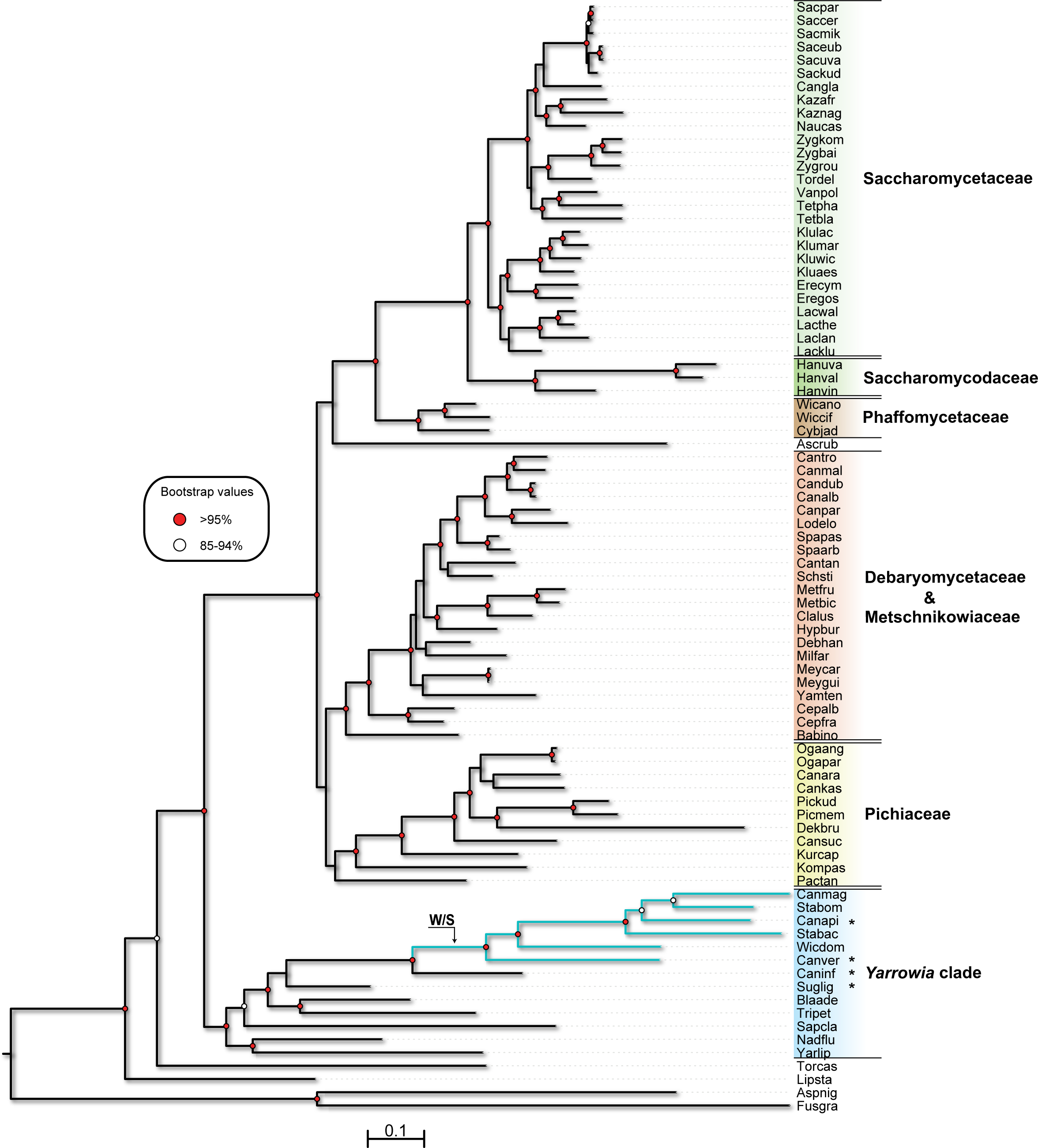
ML phylogeny of Saccharomycotina species. ML phylogeny of Saccharomycotina species. Species represented are the same as in Gonçalves et al. 2016 except for the addition of W/S-clade species C. apicola NRRL Y-50540 (Canapi*) and C. versatilis JCM 9598 (Canver*), C. infanticola DS-02 (Caninf*), and Sugiyamaella lignohabitans CBS 10342 (Suglig*). Rpa1-Rpc2 protein sequences for each species were used to construct the ML tree as in Gonçalves et al. 2016 with RAxML (Stamatakis, 2006) v7.2.8 using the PROTGAMMAILG model of amino acid substitution. Aspergillus niger (Aspnig) and Fusarium graminearum (Fusgra) were used as outgroups. Only strong branch support values (>85%) are displayed. Branches in blue represent W/S-clade species. This tree is in agreement with the recently published phylogeny of 86 yeast species based on genome sequences (Shen & Zhou, 2016).

**Figure 3 - figure supplement 1.**
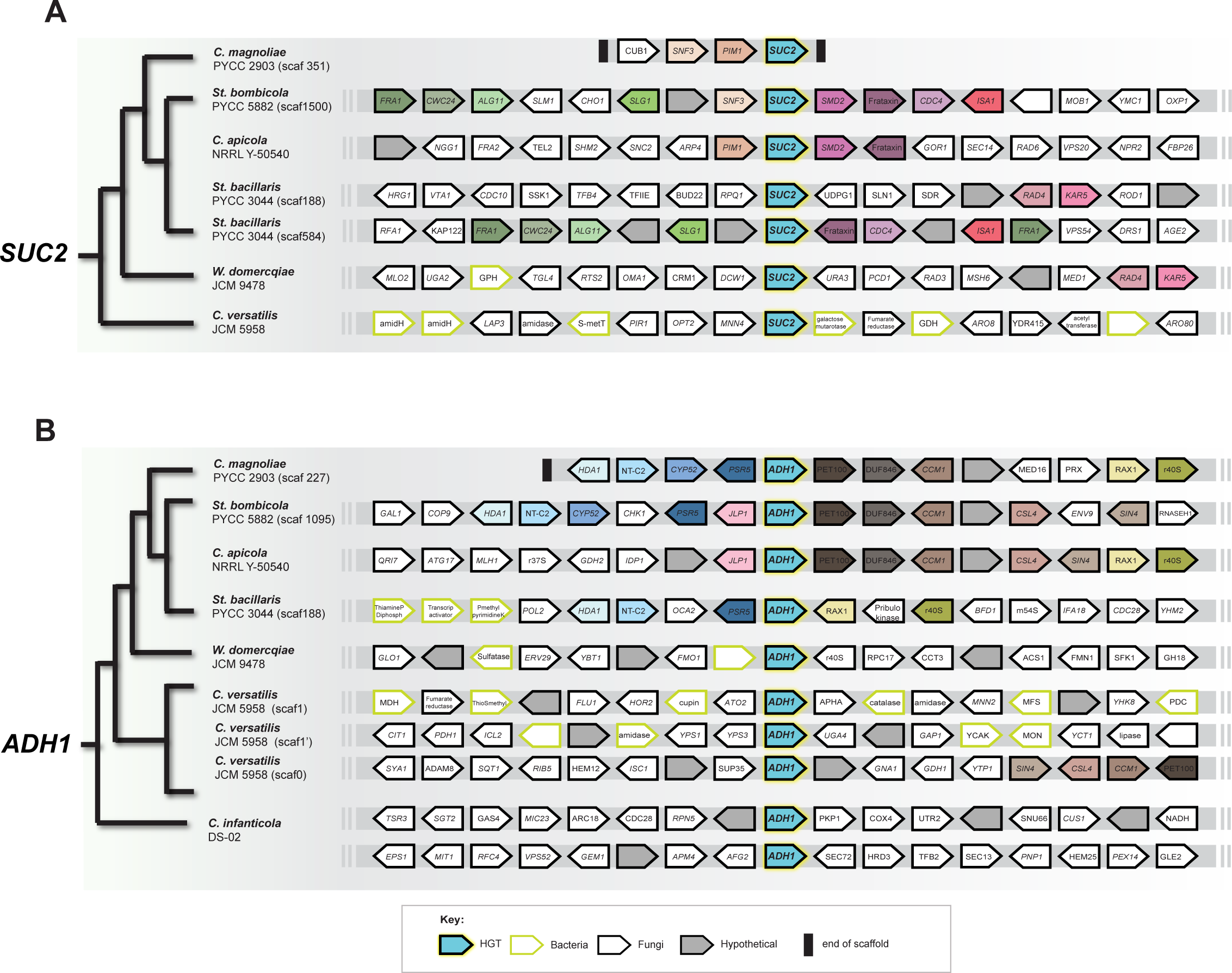
Gene content and organization in the *SUC2* (A) and *ADH1* (B) loci in W/S-clade species and C. *infanticola*. Chromosomal regions encompassing the ADH1 and SUC2 genes are depicted by grey bars for the species represented in the schematic tree on the left, which is based on the species relationships depicted in the species phylogeny (Figure 3-figure supplement 1). Horizontally transferred ADH1 and SUC2 genes are depicted as blue arrows in all species, denoting transcriptional orientation. Orthologous genes among the species examined exhibit the same color. Non-syntenic genes are colored in white, while hypothetical genes are colored in grey. Genes of apparent bacterial origin (based on top 100 BLAST hits in NCBI database) are outlined in light green and genes of apparent fungal origin are outlined in black. The end of a chromosome/scaffold is indicated by a solid black bar (as shown for C. magnoliae). Genomic loci are shown for each gene as they appear in their respective genome databases. The W. domercqiae JCM 9478 (the same strain as PYCC 3067) genome was used instead of the draft genome reported in Goncalves et al, 2016 because of its higher assembly quality.

**Figure 3 - figure supplement 2.**
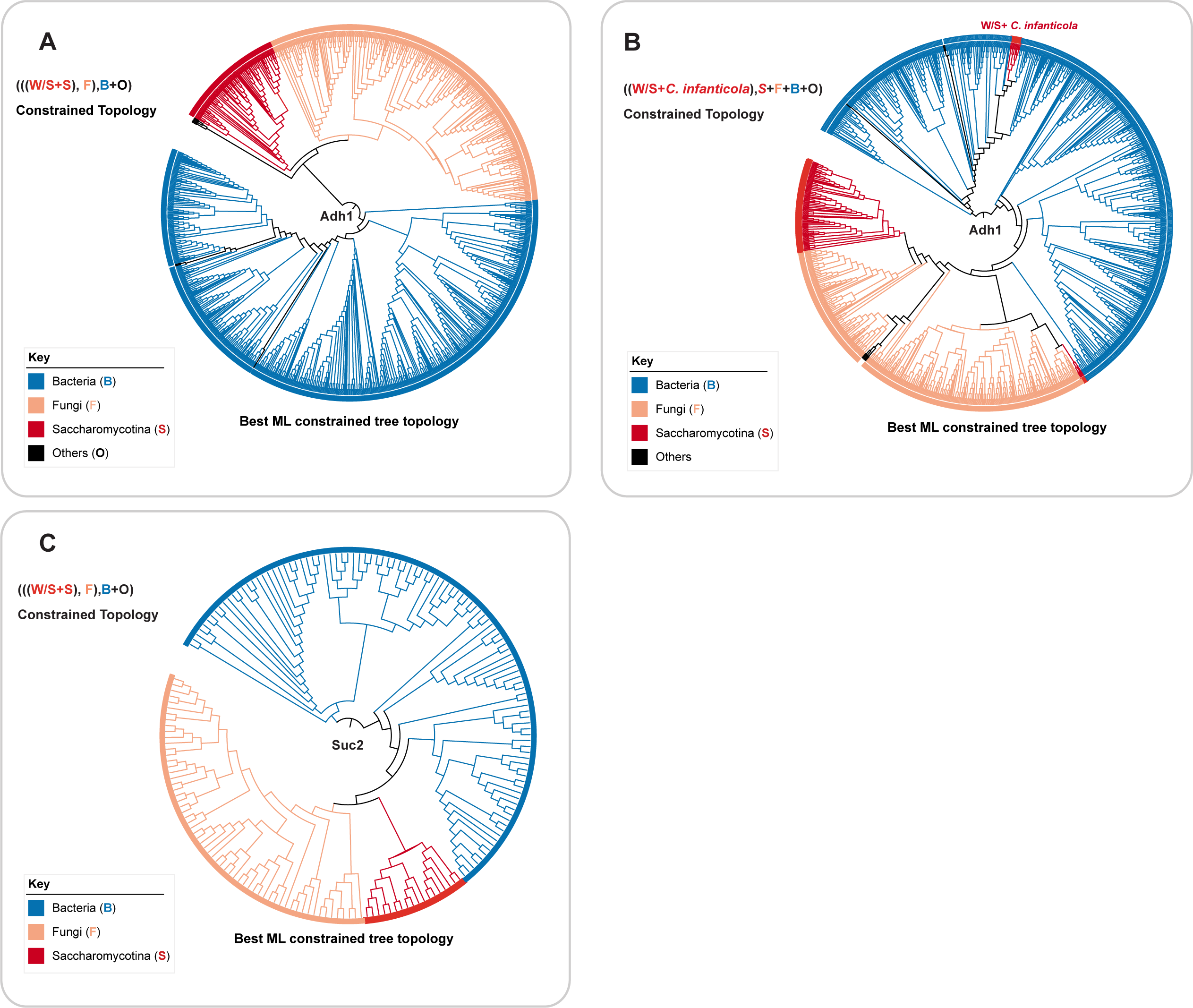
Topology test analyses for *ADH1* and *SUC2*. A) Constrained topology and best ML constrained tree for ADH1 to test HGT from bacteria. B) Constrained topology and best ML constrained tree for ADH1 to test independence of HGT events in W/S clade and C. infanticola. C) Constrained topology and best ML constrained tree for SUC2 to test HGT from bacteria.

**Figure 5- figure supplement 1.**
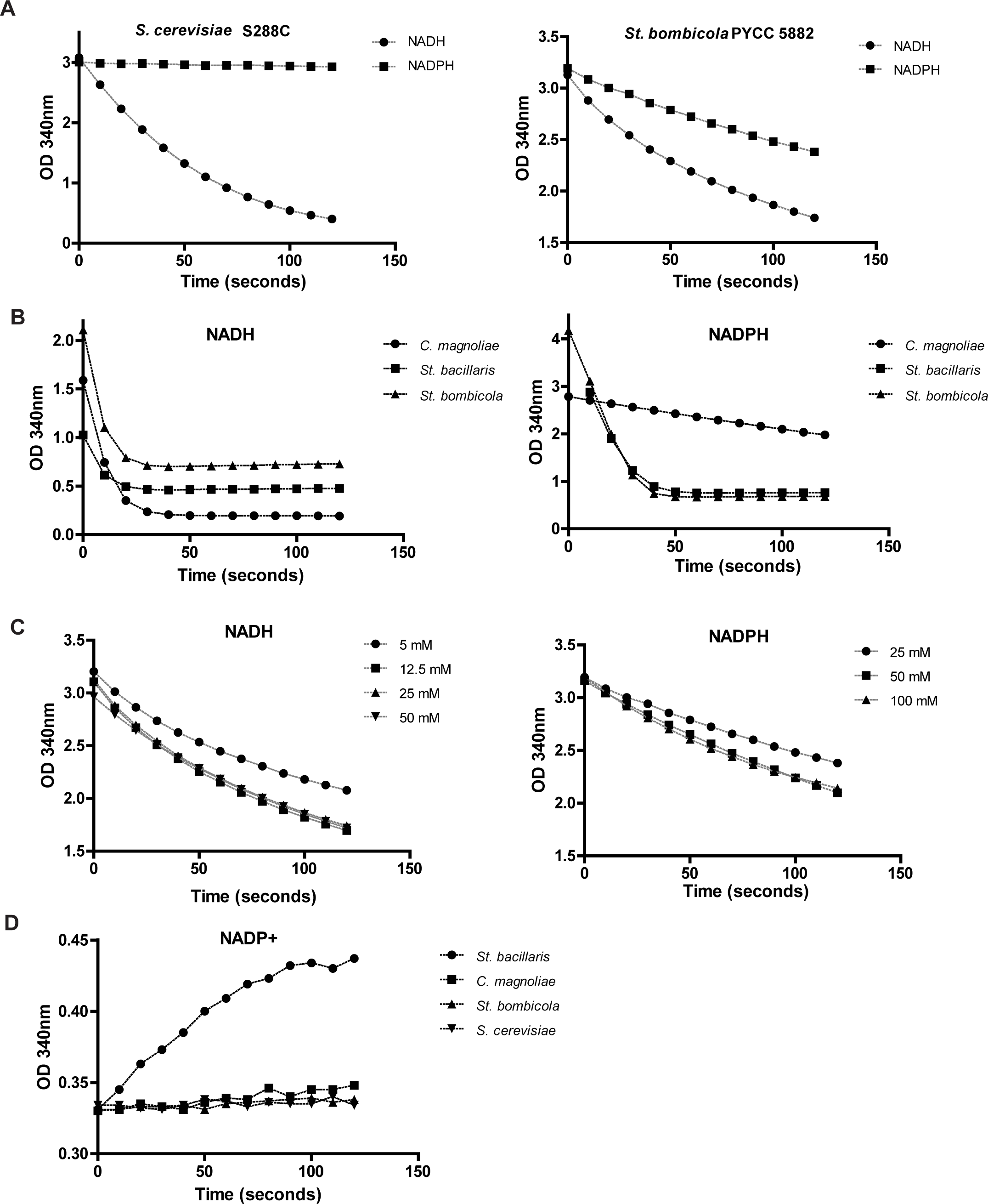
Alcohol dehydrogenase (Adh) activities. A) Adh activity in cell-free extracts of *S. cerevisiae* S2288C and *St. bombicola* PYCC 5882 cultures grown in YPD medium. B) Adh activities in cell-free extracts of other W/S-clade species when NADH or NADPH were used as cofactor. C) Adh activities in St. bombicola cultivated in 20FG medium using different concentrations of substrate (acetaldehyde) and NADH or NADPH as cofactors. Unless otherwise indicated, 100 mM of acetaldehyde and 1 mM of either NADH or NADPH were used in all experiments. D) NADP^+^-dependent glycerol dehydrogenase activity in cell-free extracts of *St. bombicola, C. magnoliae, S. bacillaris, and S. cerevisiae*

**Figure 5- figure supplement 2.**
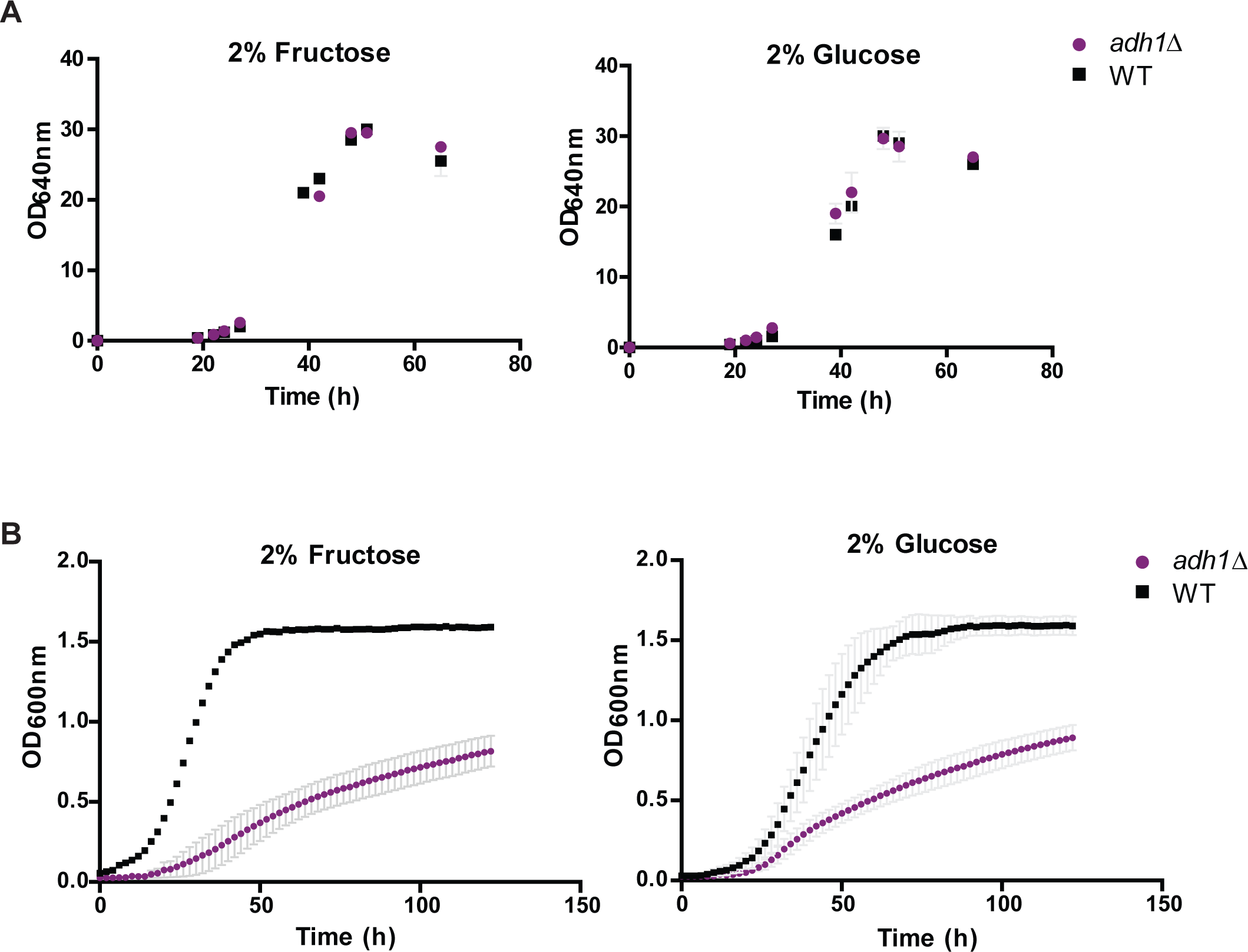
Growth of WT and adh1∆ under aeration (A) and microaeration (B). All strains were grown in minimal media supplemented with 2% (w/v) fructose or 2% (w/v) fructose. Error bars represent standard deviation of assays performed in duplicate in two biological replicates. For WT grown in fructose under microaeration (B), only one representative curve is shown for the WT strain because stochastic variation of the lag phase was observed under this condition.

**Figure 6-figure supplement 1.**
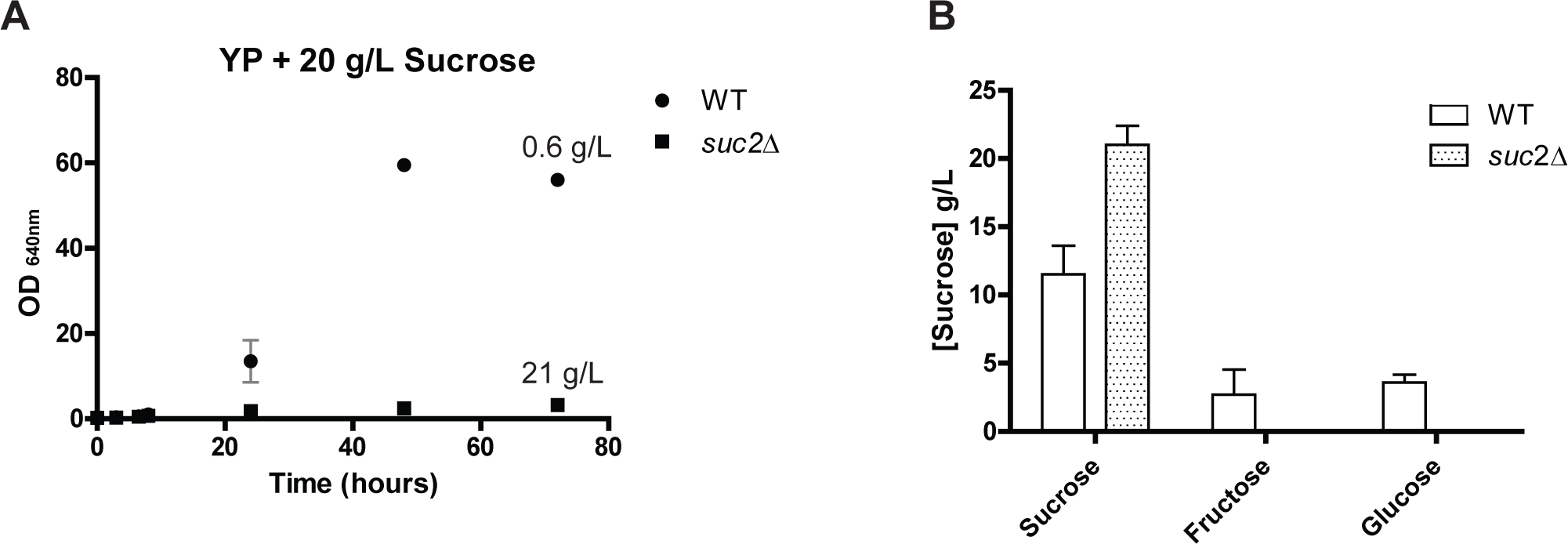
Growth and consumption of sucrose in *St. bombicola* WT and *suc2∆* mutant. A) Strains were grown in YP supplemented with 20 g/L of sucrose for 72 hours at 25°C. Sucrose concentrations determined by HPLC after 72 hours of growth were 0.6 g/L for the WT strain and 21 g/L for *suc2∆* strain. B) HPLC analysis of extracellular sugar concentrations after 24 hours of growth. Error bars represent standard deviation of assays performed in duplicate in two biological replicates.

## Source data

Figure 1- source data 1- Numerical values for glucose and fructose concentration used for the construction of Figure 1 plots.

Figure 2-source data 1- Comparative analysis of central carbon metabolism-related genes in the W/S clade.

Figure 3-source data 1- KEGG, Interpro and GO annotations of genes of bacterial origin in the W/S clade.

Figure 5- source data 1- Numerical values for the construction of plots displayed in Figure 5 and statistically significant differences between WT and deletion mutants for metabolite production and sugar consumption.

Figure 5- source data 2- Primers for construction of *St. bombicola* deletion mutants.

